# Revealing reassortment in influenza A viruses with TreeSort

**DOI:** 10.1101/2024.11.15.623781

**Authors:** Alexey Markin, Catherine A. Macken, Amy L. Baker, Tavis K. Anderson

## Abstract

Reassortment among influenza A viruses (IAV) facilitates genomic evolution and has been associated with interspecies transmission and pandemics. We introduce a novel tool called TreeSort that accurately identifies recent and ancestral reassortment events on datasets with thousands of IAV whole genomes. The algorithm has immediate relevance to modern large-scale surveillance studies that generate thousands of whole genomes. TreeSort uses the phylogeny of a selected IAV segment as a reference and finds the branches on the phylogeny where reassortment has occurred with high probability by incorporating information from other gene segments. The tool reports the particular gene segments that were involved in reassortment and how different they are from prior gene pairings. Using TreeSort, we studied reassortment patterns of different IAV subtypes isolated in avian, swine, and human hosts. Avian IAV demonstrated more reassortment than human and swine IAV, with the avian H7 subtype displaying the most frequent reassortment. Reassortment in the swine and human H3 subtypes was more frequent than in the swine and human H1 subtypes, respectively. The highly pathogenic avian influenza H5N1 clade 2.3.4.4b had elevated reassortment rates in the 2020-2023 period; however, the surface protein-encoding genes (HA, NA, and MP) co-evolved together with almost no reassortment among these genes. We observed similar co-evolutionary patterns with very low rates of reassortment among the surface proteins for the human H1 and H3 lineages, suggesting that strong co-evolution and preferential pairings among surface proteins are a consequence of high viral fitness. Our algorithm enables real-time tracking of IAV reassortment within and across different hosts, can be applied to determine how specific genes evolve in response to reassortment events, and can identify novel viruses for pandemic risk assessment. TreeSort is available at https://github.com/flu-crew/TreeSort.

## 1 Introduction

Reassortment is an evolutionary mechanism of segmented viruses, such as influenza A virus (IAV), that generates new gene combinations with the potential to exhibit novel phenotypes [48, 25]. A successful reassortment event produces viral progeny with mixed gene combinations from two or more parental viruses [31]. In IAV, there are 16 hemagglutinin (HA) subtypes and 11 neuraminidase (NA) subtypes detected in a range of wild bird hosts [54]. Though there are fewer subtypes detected in other hosts, e.g., human and swine IAV are currently restricted to H1N1, H1N2, and H3N2 subtypes, reassortment between and within subtypes constantly creates new genotypes that can affect evolutionary trajectories and phenotypes [51, 15, 52]. More broadly, the process of reassortment is critical to quantify: pandemic IAV emerged in 1957 with H2N2, 1968 with H3N2, and 2009 with H1N1pdm09 [24, 47, 32] as the result of reassortment in non-human hosts.

Advances in genomic surveillance have resulted in more than 210, 000 IAV whole genome sequences available in the public domain from human, swine, and avian hosts, with approximately 20, 000 new genomic sequences each year [46]. These data provide a unique opportunity to trace the genomic evolution of IAV, identify major reassortment events, and determine how reassortment may impact the propensity for IAV to cross host species boundaries. However, despite many previous attempts to tackle reassortment inference [41, 58, 34, 30, 33, 28, 4], there are no existing approaches that can accurately infer reassortment on datasets with thousands or tens of thousands of whole genome sequences.

To overcome this, we introduce TreeSort, a fast and accurate method that uses a rigorous statistical framework to identify both recent and ancestral reassortment events. TreeSort leverages the molecular clock signal in the evolution of individual IAV segments to accurately identify reassortment along the branches of a user-selected segment phylogeny (see Fig. 1A-B). For example, by selecting the evolution of the hemagglutinin (HA) gene as the backbone, TreeSort can identify putative reassortment events between the HA and all other gene segments. Additionally, TreeSort estimates the underlying *reassortment rate* for a given IAV dataset. The reassortment rate is the expected number of reassortment events per year as we trace back the ancestral lineage of a single sampled virus strain. In simulation studies, we demonstrated that the accuracy of TreeSort is higher than 90% on average, and it increased with an increase in the sampling density (Fig. 1C). We compared TreeSort against a Bayesian method for the joint inference of evolutionary history of all gene segments, CoalRe [33], and demonstrated that TreeSort is more accurate on simulated data, particularly on large datasets with high sampling density (median branch length of less than 3 substitutions per branch).

**Figure 1:**
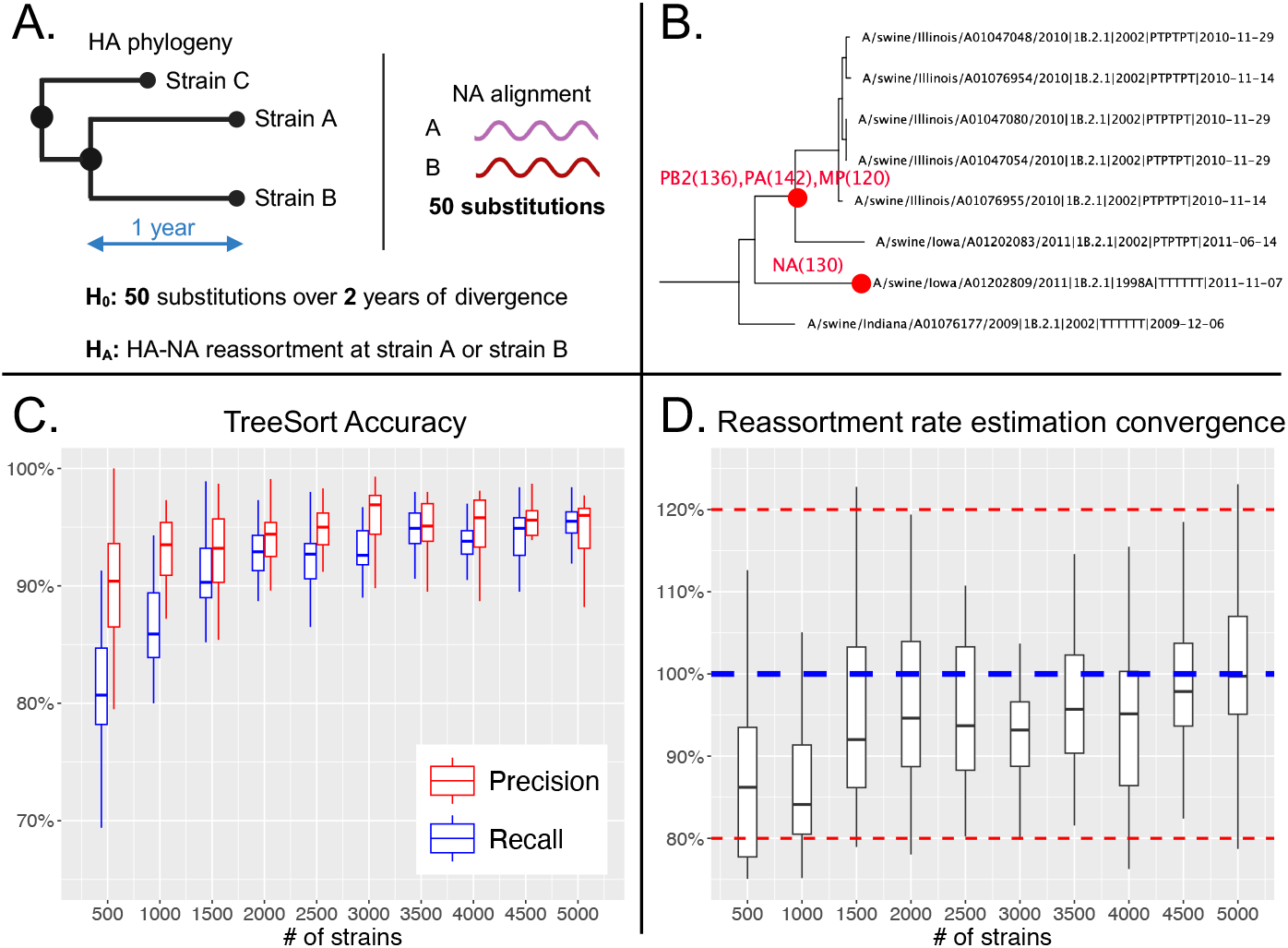
TreeSort overview and accuracy in reassortment history reconstruction and reassortment rate estimation. **A:** TreeSort uses a statistical hypothesis test to infer reassortment events along the branches of a fixed gene phylogeny. **B:** An example of output provided by TreeSort: the red circles indicate the tips of branches where reassortment has occurred, and the annotations indicate the new gene segments that the virus acquired on that branch. e.g., PB2(136) indicates that a new PB2 segment was acquired and was at least 136 nucleotides different from the ancestral PB2 prior to reassortment. **C:** The precision and recall of TreeSort increases with the increase in the sampling density in the data. Precision indicates whether the reassortment events reported by TreeSort were accurate, and recall indicates how many of the reassortment events detected by TreeSort were true. **D:** The accuracy of reassortment rate estimation by TreeSort relative to the true rate. The average and median estimated reassortment rates converged within 10% of the true rate for higher-density datasets with over 1,500 strains.

Using TreeSort’s ability to accurately estimate reassortment rates on different datasets, we compared reassortment rates across the major avian, swine, and human IAV HA subtypes. We found that avian IAV generally displayed higher reassortment rates than swine and human IAV, with the H7Nx and H5Nx avian lineages undergoing more reassortment than any other lineage from our analysis. We also conducted an analysis of reassortment in the highly pathogenic avian influenza (HPAI) H5Nx clade 2.3.4.4b viruses. This clade has recently undergone global dissemination and is associated with numerous spillovers into mammals [1, 9, 40, 20, 36].We demonstrate that this clade underwent very high rates of reassortment, approximately three reassortment events over four years for each individual strain, between 2020 and 2023. Most of the reassortment events involved the PB2 and/or the NP gene segments, and there was almost no reassortment between the HA and the NA or MP segments, suggesting that there were strong evolutionary pressures for the surface proteins to co-evolve together throughout this global outbreak.

## 2 Results

### 2.1 TreeSort demonstrated high accuracy in reassortment inference

TreeSort showed high accuracy in simulation studies with a credible model of influenza A virus evolution (Fig. 1C). Accuracy increased with an increase in the sampling density of the genomic data with precision close to or exceeding 95% on datasets with 2,000 or more strains (median branch length below 2 substitutions per branch). Note that the median branch length reflects the sampling density, i.e., lower branch lengths indicate higher sampling density (Supplemental Fig. S1).TreeSort’s recall, i.e., the percentage of true reassortment events recovered by the tool, increased from approximately 80% on datasets with 500 strains to 95% on datasets with 5,000 strains. Additionally, we compared the performance of TreeSort with CoalRe – a Bayesian reassortment network inference approach implemented in BEAST 2 [33]. When provided with a true phylogenetic tree, TreeSort was more accurate than CoalRe across all datasets (Supplemental Fig. S2). The difference was particularly noticeable on datasets with 5,000 strains when the performance of CoalRe dropped to only 40% accuracy due to the lack of convergence in the Bayesian estimation.

We also studied the ability of TreeSort to estimate the *rate of reassortment* : specifically, the tool quantifies the frequency of reassortment in the dataset per ancestral lineage per year (see Methods for the formal definition^∗^). TreeSort recovered the true reassortment rate within the 10% error margin on datasets with 1,500 or more strains and within the 5% error margin on datasets with 3,500 or more strains (Fig. 1D). The average reassortment rate estimated across multiple replicates by TreeSort was within 20% of the true rate under all sampling density regimes. In practice, we recommend estimating the reassortment rate multiple times and taking the average across replicates for more robust rate estimation.

### 2.2. Avian IAV had a higher reassortment rate relative to human and swine IAVs

We applied TreeSort to estimate reassortment rates in different HA subtypes endemic to avian, swine, and human hosts using public whole genome data collected between 2010 and 2023. Avian IAV generally had more reassortment than human and swine lineages, with the avian H7 subtype displaying the highest reassortment rates among the compared HA subtypes (Fig. 2A). The avian H7 subtype had around 0.6 reassortment events per ancestral lineage per year, almost 2-fold higher than any other HA subtype analyzed in our study. Avian H5 displayed the next highest reassortment rate of 0.3 events per ancestral lineage per year. The reassortment rates in the swine subtypes were higher than those in the human subtypes – 0.1 average rate across the swine H1 and H3 and 0.07 across the human H1 and H3. Notably, reassortment in the swine H3 subtype was more frequent than in the swine H1 subtype. Similarly, there was more reassortment in the human H3 subtype when compared to human H1s, consistent with previous studies [33]. However, the difference between the human H3 and H1 lineages was not significant: a reassortment rate of 0.08 for H3 compared to 0.07 for the H1. Note that a reassortment rate of, e.g., 0.5 corresponds to 1 reassortment event every 2 years for each individual strain (on average).

**Figure 2:**
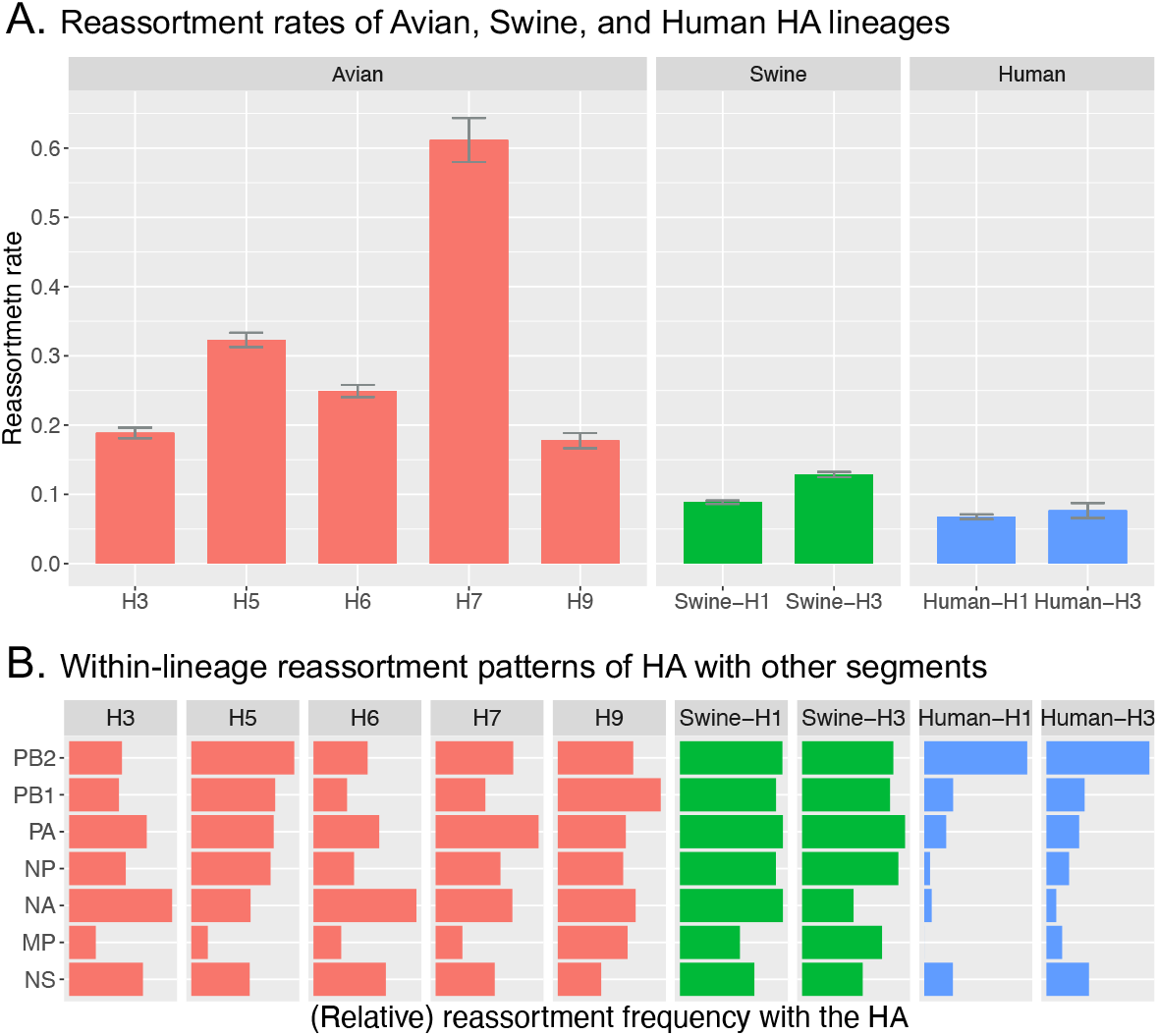
Influenza A virus reassortment patterns by host and HA subtype. **A**: Reassortment rates were expressed as the expected number of reassortment events per ancestral lineage per year and estimated by TreeSort on whole-genome data for each host and HA subtype. Here, e.g., ‘H5’ implies analysis over the H5Nx genomes. A reassortment rate of 0.5 suggests that a single virus has roughly a 50% probability of reassortment, on average, over the course of a year. **B**: Relative reassortment rates between the HA and the other gene segments for each lineage. The rates were normalized to be between 0 and 1, with 1 indicating the highest reassortment rate across the 7 gene segments.

### 2.3 Reassortment patterns and preferential gene pairings varied between hosts and subtypes

Linkage between different gene segments can affect how frequently reassortment occurs between those segments. Here, we studied the reassortment patterns relative to the HA gene across the avian, swine, and human HA subtypes (Fig. 2B). For most of the subtypes, the MP segments reassorted least relative to the HA, suggesting evolutionary constraints on these two genes to co-evolve. For avian-H5, swine-H3, human-H1, and human-H3, we also demonstrate relatively low reassortment rates of NA relative to the HA, suggesting that HA-NA balance plays an important role in the evolution of these surface proteins as previously reported [8, 12]. The HA-NA reassortment rates were particularly low in the human IAV subtypes. In contrast, the avian-H3 and avian-H6 lineages had very high rates of HA-NA reassortment, suggesting that the HA-NA balance may not be as important to the fitness and evolution of IAV in those subtypes. Generally, the HA-PB2 pair was the most permissive for reassortment, and the reassortment of PB2 relative to a fixed HA was particularly frequent in the avian-H5 and human IAVs.

### 2.4 Changes in reassortment patterns of the H5 clade 2.3.4.4b avian IAV after 2020

The H5 2.3.4.4b avian HPAI clade was first detected around 2010, and circulated in Eurasia primarily paired with N6 or N8 neuraminidase. In late 2020, a novel H5N1 2.3.4.4b virus emerged and disseminated globally in a panzootic outbreak [21, 57]. This shift to N1 in 2020 and a subsequent panzootic have occurred in conjunction with an increased reassortment rate – 0.72 reassortment events per ancestral lineage per year after 2020 and 0.44 prior to 2020 (Fig. 3). Additionally, there were differences in the number of nucleotide changes induced by the reassortment events. During the 2020-2023 period, the genetic divergence between reassorting PB2, PB1, PA, and NP segments and the segments that they replaced was greater than the genetic divergence of those reassorting in the prior period (2013 - 2020) (see Supplemental Fig. S3). This shift is likely due to the spread of the virus to the Americas in 2021 [7, 16] and extensive reassortment with the local low pathogenic avian influenza viruses [57]. Additionally, the post-2020 period was associated with increased substitution rates across all eight segments (Supplemental Table S1).

**Figure 3:**
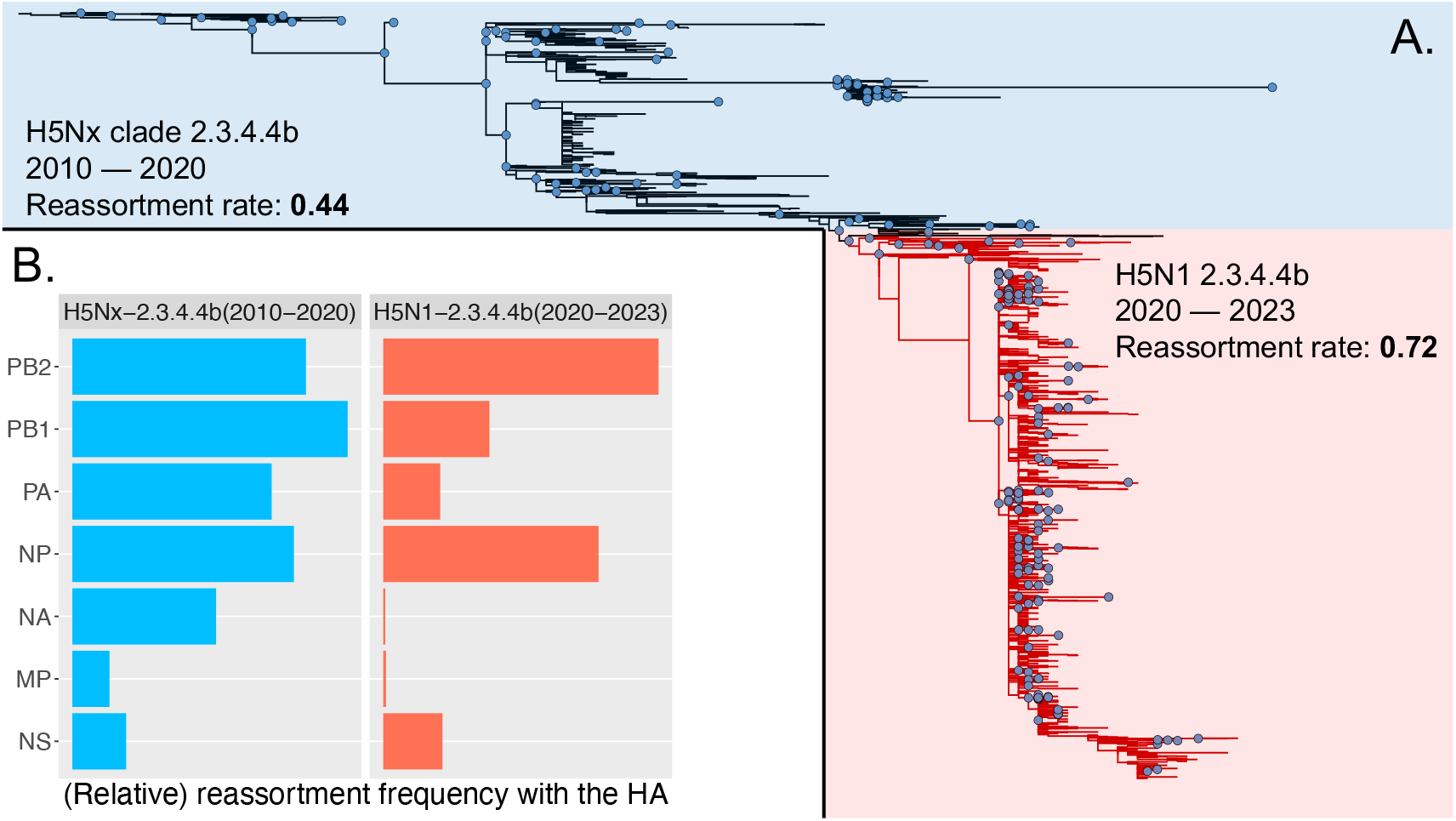
The HA phylogeny split into two parts (A) and a comparison of the reassortment rates and reassortment patterns (B) in the HPAI H5Nx clade 2.3.4.4b before the N1 reassortment in late 2020 and global dissemination (blue) and after this major reassortment event (red). The nodes on the phylogeny highlighted with blue circles represent the inferred reassortment events.

Further, we observed a significant shift in the segment-wise reassortment frequency between the pre-2020 and the post-2020 periods (Fig. 3B). The reassortment patterns for the H5N1 2020-2023 period were similar to the human H1 and H3 patterns seen in Fig. 2B. In all three cases, the highest reassortment frequency occurred between HA and PB2, with almost no reassortment between HA and NA or between HA and MP. The major difference between the H5N1 and human IAVs was a high HA-NP reassortment frequency for the H5N1 IAVs during the period of 2020-2023. We hypothesize that low reassortment rates and strong preferential pairings among the surface protein-encoding genes, HA, NA, and MP, are indicators of high virus fitness.

## 3 Discussion

Tracking and assessment of IAV evolution in real-time is critical for animal and human health, and reassortment is a key contributor to IAV evolution. Although many reassortment events in IAV may have deleterious or neutral effects [52], others can modify transmission efficiency [51] or increase virus fitness through enabling escape from immune recognition [52, 31, 23]. Consequently, algorithms that can efficiently identify novel reassortant viruses are necessary to inform the prioritization of viral strains for phenotypic characterization. TreeSort is well suited to these analyses. It can track the evolution of IAV on the whole genome level with accuracy and efficiency (Fig. 1). Additionally, it can identify lineages with unusually high or low rates of reassortment and detect evidence of preferential gene pairings. Novel reassortant strains and genotypes identified by TreeSort can be selected for experimental investigations of the impact of reassortment on virus phenotype.

We capitalized on the computational efficiency and accuracy of TreeSort to analyze reassortment in whole genome datasets from large-scale surveillance systems of IAV in human, swine, and avian host species, and across the major endemic subtypes for each host group. Prior research has demonstrated reassortment rates that were associated with host type (wild vs. domestic) [26]. Our analyses indicate a clear host effect on the overall reassortment rate (Fig. 2). Reassortment rates in human hosts were lower than in swine hosts, which were, in turn, lower than in avian hosts. Avian hosts of IAV are highly heterogeneous, including, for example, aquatic, passerine, predatory, and poultry species. IAV from this wide range of host species provides a highly diverse substrate for reassortment; similar to a force of infection calculation, more genomic diversity leads to more reassortment. Estimates of avian IAV reassortment rates range from 0.18 to 0.61 events per ancestral lineage per year (Fig. 2A). Interestingly, the two subtypes that harbor highly pathogenic avian IAV, H5 and H7, displayed the highest reassortment rates. In an early study of reassortment of the replication complex (PB2, PB1, PA, and NP) among mixed subtypes of avian IAV, a conservative estimate of the probability that a segment would not reassort in one year was 0.95 [28], which equates to a probability 0.26 that at least one segment reassorts in a year assuming that six segments are available to reassort (omitting the possibility of MP reassorting, which is quite low in all subtypes studied here except H9). This probability is comparable to the rate of reassortment calculated here for all except H7 viruses, which exhibited the highest reassortment rate in our study.

Rates of segment-specific reassortment during the HA evolution vary across host-subtype groups (Fig. 2). Within swine hosts, these rates were comparable for all segments and consistent across H1 and H3 lineages. Within human hosts, H1 and H3 lineages of IAV had the same pattern of significantly restricted reassortment. Interestingly, although the overall reassortment rate for swine hosts is not greatly different from the reassortment rate for human hosts (Fig. 2A), the reassortment pattern by segment differed markedly between these two hosts (Fig. 2B). Human H1N1 viruses (2010 – 2023) originated from a 2009 introduction of swine H1N1 viruses. Low relative rates of reassortment of NA and MP with HA in the human H1 lineage suggest co-evolution among the surface proteins in human but not swine hosts. The step-change in dynamics may reflect the constraints on IAV evolution in the human host. Previous work has also studied the level of co-evolution among the IAV segments for human IAV [17] and swine IAV [59, 15], but on significantly smaller datasets.

Further, segment-specific reassortment rates varied substantially within and among avian lineages (Fig. 2). MP reassorted infrequently, indicating co-evolution with HA for all subtypes except H9 (Fig. 2B). Further investigation of the H5 lineage revealed a step-change in the reassortment process during the evolution of the HPAI H5 clade 2.3.4.4b viruses. The first phase (2010-2020) featured relatively unrestrained reassortment of the PB2, PB1, PA and NP segments. In the second phase (2020-2023), the PA and PB1 segments showed a reduced rate of reassortment, while the PB2 and NP segments retained high rates of reassortment. An earlier study of avian H5N1 IAV during a period of rapid evolution (2000 – 2008) identified 47 reassortment events in a much smaller dataset [35]. Of these 47 events, MP, NS, and PB1 reassorted the least (4/5 times each), followed by PB2 and PA (9/10 times each) and NP (15 times). This pattern shows similarities to that from our TreeSort analysis for the second phase of the H5Nx clade 2.3.4.4b evolution and together suggest co-adaptation pathways to achieve high fitness. The contemporary H5N1 clade 2.3.4.4b viruses caused a panzootic outbreak with frequent spillovers into mammalian hosts, including the widespread infection of dairy cattle in the US in 2024 [21, 57, 37], a highly unusual phenotype in the history of avian H5 IAV infections. Though the acquisition of new PB2 and NP segments has an unknown impact on virus phenotype, it is notable that the H5N1 2.3.4.4b virus genotype that emerged in 2024 in dairy cattle in the United States was a reassorted virus with new PB2 and NP genes – the B3.13 genotype that emerged in wild birds a few months before the spillover into cattle [37].

Strategies for surveillance of emerging variants of concern have been successfully employed based on a knowledge base of mapping from molecular change to phenotype for SARS-CoV-2 and influenza viruses [42, 13]. In contrast, to the best of our knowledge, no strategies for surveillance of novel reassortments of concern have yet been developed. Little is known about the relationship between reassortment and phenotype. The relationship is likely to be complex, as it involves multiple segments reassorting in multiple combinations. However, by analyzing large data sets, TreeSort precisely identifies many reassortment events in the process of evolution of the HA (or any other IAV segment) across host species and subtypes. This generates a rich repertoire from which to select strains for in vitro and/or in vivo experimental studies to assess how the reassorted genotype impacts phenotype (e.g., [22, 29, 27]).

Our studies also indicated two instances in which qualitative changes in the dynamics of reassortment were associated with the emergence of a dominant lineage (human H1N1pdm09 and avian HPAI H5N1). We hypothesize that the restrictions on reassortment observed in these instances of the emergence of dominant phenotypes may be indicative of the evolution of superior fitness and could provide targets for surveillance of new reassortants of concern. In both cases, the reduction in reassortment frequency of the surface protein-encoding segments (HA, NA, and MP) was of particular note. These observations provide a target for early-warning systems, where clades of viruses with the lack of HA-NA-MP reassortment can be flagged and studied for potentially enhanced transmission phenotype.

TreeSort lays the foundation for reassortment inference on very large genomic datasets. However, future studies are necessary to better understand how robust TreeSort is to variation in substitution rates across segments and possible co-evolution among segments during reassortment events. Future iterations of TreeSort will incorporate more nuanced models of evolution, such as the uncorrelated lognormal relaxed molecular clock model [10], to improve the robustness of the method to significant swings in substitution rates across the segments. Integration of such models presents a computational challenge, and novel algorithms need to be developed to make such integration efficient.

## 4 Methods

### 4.1 Preliminaries

Consider two virus strains *A* and *B* with two gene segments denoted *A*_1_, *A*_2_ and *B*_1_, *B*_2_, respectively. Let *e*_1_ denote the phylogenetic distance between *A*_1_ and *B*_1_; i.e., *e*_1_ is the expected number of substitutions per site in segment 1 between *A* and *B*. Under the Jukes-Cantor (JC69) [18] substitution model, the probability of a substitution occurring over ‘time’ *e*_1_ is

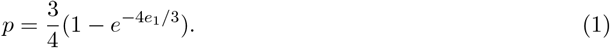

Next, let *h*_2_ denote the number of substitutions (the Hamming distance) between sequences *A*_2_ and *B*_2_. If there was no reassortment between *A* and *B, h*_2_ substitutions must have occurred over time *e*_1_. Then, under the null hypothesis of no reassortment, *h*_2_ ∼ *Binomial*(*p, n*), where *n* is the number of sites in segment 2.

A *reassortment event* is an acquisition of 1 or more novel segments (up to 7 for IAV) relative to a fixed reference segment. An *ancestral lineage* is the path from a leaf of a phylogeny (a sampled virus) to the root (most recent common ancestor), and the reassortment rate is measured in terms of the number of reassortment events per ancestral lineage per year, as defined below.

### 4.2 The TreeSort algorithm

TreeSort takes as an input the multiple-sequence alignments and inferred phylogenetic trees for each of the gene segments involved in the reassortment analysis. One of the segments, e.g., the HA gene, must be used as a reference phylogeny, and the reassortment events will be mapped on the branches of that phylogeny. Note that the reference phylogeny needs to be rooted, which may be achieved through software such as TempEst [43] or TreeTime [45]: other trees may be unrooted.

#### 4.2.1 Estimating substitution rates

TreeSort estimates the substitution rates (number of substitutions per site per year) for each input gene segment using the TreeTime Python package [45]. We use TreeTime’s functions of finding an optimal root under the autocorrelated molecular clock model and JC69 nucleotide substitution model. TreeTime detects and ignores molecular clock outliers. For large datasets with over 1,000 taxa, we reduced computational time for evolutionary rate estimation by randomly subsampling 1,000 taxa for each segment independently and performing the TreeTime analyses on the subsampled data.

We denote an inferred evolutionary rate for segment *S* by *r*_*S*_.

#### 4.2.2 Dynamic algorithm for reassortment inference

Consider two gene segments *R* and *Q*, where *R* is the *reference* segment and *Q* is the *query* segment. Our goal is to map reassortment events between these two segments on a phylogeny of the reference segment *R*.

Let ⟨*R*⟩ = *R*_1_, …, *R*_*n*_ and ⟨*Q*⟩ = *Q*_1_, …, *Q*_*n*_ denote (aligned) sequences for segments *R* and *Q* for *n* different strains, and *T*_*R*_ denote a rooted maximum-likelihood phylogeny for segment *R*. We solve the small parsimony problem on a tree *T*_*R*_ relative to the alignment ⟨*Q*⟩ using Fitch’s algorithm [11] as implemented in DendroPy 4 [49]. That is, for every pair of sibling nodes *u, v* in *T*_*R*_, we determine the smallest number of substitutions that are required to explain the differences between sequences in the subtrees rooted at *u* and *v* in *T*_*R*_, assuming that ⟨*Q*⟩ evolved according to tree *T*_*R*_ in those subtrees. Additionally, if *u* and *v* have an aunt node *x* (Figure 4A), we compute the parsimony distance between *u* and *x* and between *v* and *x*. That is, we obtain a parsimony distance for each pair of sibling nodes (*u, v*) and for each node-aunt pair (*u, x*). We denote the parsimony distance over segment *Q* between two nodes as *d*_*Q*_(*u, v*) and the original maximum likelihood distance in tree *T*_*R*_ by *d*_*R*_(*u, v*).

**Figure 4:**
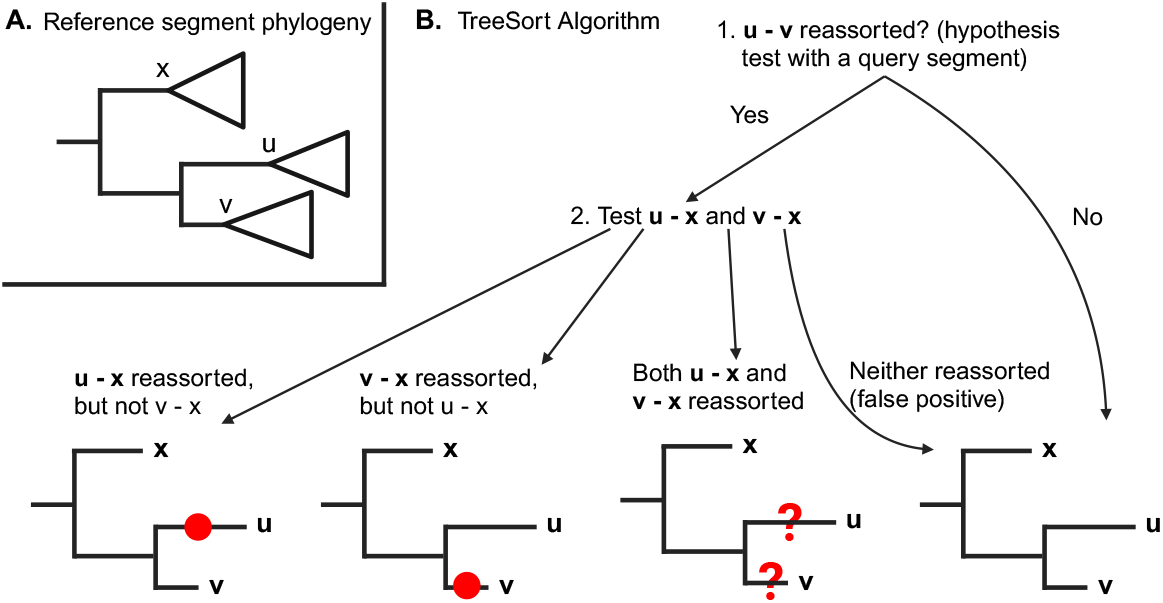
**A**: Example of two sibling nodes *u* and *v* and their ‘aunt’ node *x* on tree *T*_*R*_. **B**: TreeSort uses an aunt node/clade (node *x*) to determine the branch of the tree where the reassortment event has happened between u and v. The red circles indicate the inferred reassortment placement and question marks indicate insufficient resolution to confidently place the reassortment event onto one of the two branches. That is, the immediate neighbors on the phylogenetic tree may be too distant or have undergone reassortment as well. Recall that, e.g., *u* − *x reassorted* is a shortcut for “there is a reassortment event on the path between *u* and *x*”.

We perform a clock-based hypothesis test to determine whether reassortment has occurred on the path between *u* and *v*. Note that if there is no reassortment on the path connecting these two nodes, then segments *R* and *Q* have evolved over the same period of time, and the expected number of substitutions per site for segment *Q* is 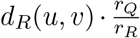, 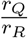 where is the ratio of substitution rates between these segments.

#### 4.2.3 Reassortment test between nodes *u* and *v* of *T*_*R*_

The following hypothesis test allows us to check for reassortment between two nodes. *H*_0_(*u, v*): *no reassortment on the path between u and v. That is*,

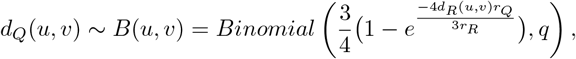

*where q is the length of the alignment* ⟨*Q*⟩.

*H*_*A*_(*u, v*): *there was a reassortment event on the path between nodes u and v involving an acquisition of a new segment Q*.

We reject the null hypothesis when *d*_*Q*_(*u, v*) is in the right tail of the *B*(*u, v*) distribution with a p-value cutoff of *α* = 0.001. Note that this is a default cutoff value that the user can adjust. For simplicity, we say that *u* − *v reassorted* if *H*_0_(*u, v*) is rejected.

The algorithm proceeds bottom-up on the reference tree *T*_*R*_ to examine each pair of sibling nodes *u, v* and uses the above hypothesis test to identify reassortment events as illustrated in Fig. 4B. For every pair of sibling nodes *u, v*, and the respective aunt node *x* (if exists), the algorithm checks if *u* − *v* reassorted, and in that case, checks if *u* − *x* and/or *v* − *x* reassorted. Note that the algorithm determines either a certain reassortment placement or an uncertain placement as denoted by question marks in Fig. 4B. In case of a certain placement, the inferred edge gets the “query(*d*_*Q*_(*u, v*))” annotation, e.g., “PB2(120)”, where the number in the parentheses denotes the lower bound on the number of substitutions between the old and the new query gene segments post-reassortment. In the case of uncertain placement, both sibling edges get the “?query(*d*_*Q*_(*u, v*))” annotation, e.g., “?NP(50)”. TreeSort processes each query segment independently and then combines the inferred reassortment annotations for each of the edges of the reference phylogeny.

Note that for a dataset with *n* genomes, TreeSort implements the decision tree in Fig. 4B *n* − 1 times for each query segment. Therefore, to control the false positive rate for reassortment detection of a segment, *α*_0_, one can set the p-value in TreeSort to be *α*_0_*/n*. E.g., for *α*_0_ = 0.01 (1%) and *n* = 1000, p-value cutoff should be set to 0.00001.

#### 4.2.4 Resolving multifurcations for most parsimonious reassortment inference

TreeSort will help resolve multifurcations in the reference tree *T*_*R*_ using the information from the other gene segments. This procedure ensures that TreeSort does not treat related reassortment events as independent events and results in more consistent reassortment rate estimates. To achieve this, prior to the gene-by-gene reassortment inference procedure outlined above, TreeSort concatenates all the query (non-reference) gene segments into a single alignment *Q*_*all*_ and computes the averaged substitution rate *r*_*all*_ over those segments taking into account the length of the query gene segments. TreeSort then solves the small parsimony problem on *T*_*R*_ using the concatenated alignment. For every node *v* with more than two children in *T*_*R*_, TreeSort performs the following procedure:

1. Randomly order the children of *v*.
2. Bin the child nodes into non-reassorted subtrees using the above reassortment test and *r*_*all*_ as the query substitution rate. In particular, those non-reassorted subtrees are constructed progressively over the ordered child nodes, with new children being placed on top of the appropriate subtree and the Fitch parsimony inference propagating to the new root of that subtree.
3. Merge the formed non-reassorted subtrees into a single caterpillar-tree structure and attach back to *v*.

#### 4.2.5 Accounting for the deviation from the strict molecular clock

As IAV segment evolution does not always adhere to a strict molecular clock (e.g., [55]), we extend our hypothesis test from above to relax the clock constraints. We introduce a clock-deviation parameter *λ* ≥ 1, such that the substitution rate of a segment *S* on a particular tree branch *e* can vary in the 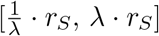, *λ r*_*S*_] interval, where *r*_*S*_ is the base rate estimated above. Then, in the extreme case for the *H*_0_(*u, v*) hypothesis testing, the substitution rate along the (*u, v*) path in segment *R* may be 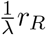 and in segment *Q, λr*_*Q*_. That is, the maximum substitution rate for segment *Q* is 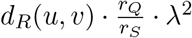. Then,

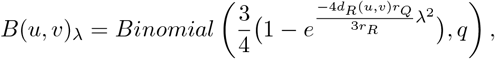

and we adjust our hypothesis test accordingly.

### 4.3 Estimating the clock-deviation parameter *λ*

We used the avian influenza A substitution rate analysis for H3N8 and H7N7 subtypes by Worobey et al. [55] to estimate the parameter *λ*. We analyzed the maximum clade credibility (MCC) trees for the PB2, PB1, PA, H3, H7, NP, N7, N8, MP, NS gene segments constructed by Worobey et al. using the uncorrelated lognormal distribution (UCLD) relaxed clock model [10] that allows for an uncorrelated branch-specific substitution rate. For each tree, we computed the median, 5th percentile, and 95th percentile substitution rates among the tree branches. For simplicity, we refer to the 5th percentile rate as the min rate and the 95th percentile rate as the max rate. Table 1 summarizes the deviation of the substitution rates from the median rate towards the min and the max for each gene segment. For six out of ten segments, the deviation ratio did not exceed 2, and the maximum overall deviation was 2.45. Therefore, by default, TreeSort uses *λ* = 2 for reassortment inference, but this parameter can be adjusted depending on the molecular clock signal in the dataset.

**Table 1:** Substitution rates deviation from the median rate for each gene segment of avian influenza A subtypes H3N8 and H7N7 estimated from MCC trees in [55].

| Segment | Median-min ratio | Max-median ratio |
| --- | --- | --- |
| PB2 | 1.76 | 1.95 |
| PB1 | 1.59 | 1.64 |
| PA | 1.65 | 1.72 |
| H3 | 1.84 | 2.07 |
| H7 | 2.08 | 1.87 |
| NP | 1.71 | 1.88 |
| N7 | 1.31 | 1.35 |
| N8 | 1.83 | 2.17 |
| MP | 1.57 | 2.13 |
| NS | 1.88 | 2.45 |

### 4.4 Estimating the reassortment rate

TreeSort processes all non-reference segments and collates the reassortment annotations onto the branches of the reference tree. These data are used to estimate the rate of reassortment.

We model reassortment as a Poisson point process, starting at the root of the reference phylogeny *T*_*R*_, assumed to be bifurcating, and going to the leaves of *T*_*R*_, duplicating at every branching point (node). This is similar to the modeling of gene duplications and gene losses in phylogenomics [3, 44]. Here, we interpret reassortment to mean the exchange, at a single point in time, of 1 or more segments in a genome constellation. We define *ρ* to be the rate, in events/year, of this reassortment process.

Note that TreeSort cannot identify the donor of reassorting segments. Hence, when multiple annotations such as “PB2(120),NP(105)” are assigned to a single branch of *T*_*R*_, it is not possible to know if PB2 and NP reassorted as a single event or as multiple consecutive events. In densely sampled surveillance studies, when branch lengths of the phylogenies are short, we believe that multiple consecutive events on a single branch are unlikely, and that branches with more than one annotation reflect a single reassortment event. With decreasing density of sampling, the likelihood that multiple annotations arise from consecutive events increases. We provide a numerical estimator of the rate of reassortment applicable to all datasets, 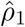, and a simple formula for an approximation to the rate that is accurate in the case of high-density sampling, 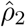.

To derive 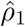, we score branches in the reference tree according to whether or not the branch is annotated. For branch *e* of *T*_*R*_, let *B*_*e*_ = 1 if it is annotated and 0 otherwise. Under the Poisson model, the probability of no reassortment events along a branch of length *l*_*e*_, measured in years, is 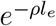, and the probability of at least one reassortment event is 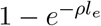. Then the likelihood function is

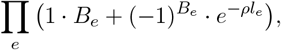

and we can use numerical optimization to maximize this function with respect to *ρ* to obtain 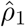. In practice, the estimator needs to be modified to account for uncertain annotations, e.g., “?PA(56)”. Branches with at least one certain annotation are assigned *B*_*e*_ = 1. For a branch *e* with only uncertain annotations, we identify its sister branch *e*^′^, and include the probability that at least one event occurred over time *l*_*e*_ +*l*_*e*_*′* into the likelihood function above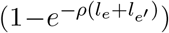. Note that we include this probability at most once per each pair of sister branches *e* and *e*^′^.

The second estimator applies to large, densely sampled datasets for which branch lengths are typically short. It assumes that multiple annotations on a single branch result from a single reassortment event. Then the expected number of reassortment events across the reference tree is, where *l*(*T*_*R*_) = Σ_*e*_ *l*_*e*_. For *B*_*e*_ = 1 if branch *e* is annotated and 0 otherwise, Σ_*e*_ *B*_*e*_ is the number of branches with at least one annotation from TreeSort. Then

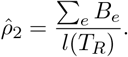

To account for uncertain annotations in the estimator 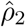, we set *B*_*e*_ = 1 if at least one annotation assigned to an edge is certain, and *B*_*e*_ = 0.5 if all annotations assigned to an edge are uncertain. In practice, datasets with high sampling density often have a few long branches that can represent historic periods with low sampling or errors in the data. To make sure that the 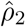 estimator does not get biased by such long branches, we remove the longest 1% of branches when computing *l*(*T*_*R*_) and Σ_*e*_ *B*_*e*_.

In simulations, we showed that 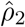 consistently falls within 10% of the true rate and converges to *ρ* as the sampling density increases (Fig. 1D).

We implemented both estimators of *ρ* in TreeSort. To find 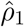, we use the Sequential Least Squares Programming (SLSQP) constrained optimization method implemented in SciPy [53] with the constraint 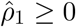. We use 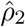 as the starting value for optimization of 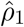.

Figures in this report use the 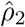 estimator, which is suitable for the high-density datasets that we analyze. For smaller and/or lower-density datasets, we recommend the 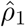 estimate. Supplemental Fig. S4 complements Fig. 2A and shows the 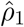 estimates for the reassortment rates across the avian, swine, and human IAV subtypes.

### 4.5 Simulating influenza A evolution with reassortment

We simulated the evolution of two IAV segments using the Coalescent with Reassortment model introduced by Muller et al. [33] and implemented in BEAST 2 [5, 6]. Similar to Muller et al., we set the reassortment rate to 0.2 events per ancestral lineage per year – this translates to the 0.1 effective reassortment rate since 50% of the time, the two segments end up in the same ancestral lineage per model definition. We sample the effective population size (EPS) parameter from a normal distribution 𝒩 (20, 0.5^2^) and the substitution rate from 𝒩 (0.005, 0.001^2^). The mean EPS corresponds to influenza A virus in swine estimates from [59], and the mean substitution rate corresponds to typical substitution rates of HA and NA genes of IAV in human and swine hosts. The sampling dates were drawn from a uniform distribution over a 20-year period, and the number of samples/strains (*n*) varied between 200 and 1000 with a step of 200 and between 500 and 5000 with a step of 500. The sequences were simulated according to the GTR+Γ substitution model [50, 56] with model parameters drawn from the distribution previously estimated for swine IAV in [30], and all sequences had a length of 1500 nucleotides – roughly corresponding to the average IAV gene length. For each sampling regime (each *n*), we simulated 25 replicates, each containing a pair of gene trees and corresponding tip sequences. The reassortment events were projected onto one of the gene trees using custom Java code that parsed the simulated phylogenetic network. As a result, for each replicate, we obtained a list of clades that appeared as a result of reassortment between the two genes.

Then, TreeSort was executed on each replicate using the simulated gene trees and sequences as the input, the p-value cutoff of 0.001, the deviation parameter of 1.1, and the option to output the list of reassorted clusters. The TreeSort reassortment events were compared to the true reassortment events, and we estimated the Precision, Recall, and F1 accuracy metrics [38]. Additionally, we compared the performance of TreeSort with CoalRe by Muller et al. [33] – a Bayesian method that estimates a reassortment network combining the evolutionary history of multiple genes while accounting for reassortment. We ran CoalRe v1.0.4 on datasets with *n* = 200, 400, 600, 800, 1000, 5000 with standard priors and setting the mean of the prior for the reassortment rate to the true value of 0.2. We set the Markov Chain Monte Carlo chain length to 5, 000, 000 samples, and the summary MCC network was obtained with a 10% burn-in rate. The Precision, Recall, and F1 metrics were computed for the resulting CoalRe estimates. To minimize the influence of smaller reassortment events with insufficient evolutionary signal for inference, we only considered the “productive” reassortment events with at least 2 years of divergence between the two parental lineages when evaluating the precision of TreeSort and CoalRe.

### 4.6 Reassortment analysis over the most prevalent avian, swine, and human subtypes

We downloaded all IAV gene sequences from the NCBI Virus database [14] with at most 20 ambiguous characters; n=971,784 [accessed on November 12, 2023]. We kept those strains that (i) were collected after 2010, (ii) had whole genomes sequenced, and (iii) were associated with the avian H3, H5, H6, H7, or H9, swine H1 or H3, and human H1 or H3 subtypes. If the resulting dataset for a given subtype contained more than 10,000 strains, we randomly sampled 10,000 strains from it. We split our analysis into the HA subtypes, aligned sequences for each segment using MAFFT v7.475 [19], trimmed alignments to the coding region, and inferred phylogenetic trees with FastTree v2.1.11 [39] for each gene segment using the GTR+Γ substitution model. For each lineage, we randomly sampled 80% of the strains in the dataset and applied TreeSort to the sampled data (10 replicates). We estimated the overall reassortment rates as outlined above, as well as the reassortment frequency between HA and each other gene segment. TreeSort was applied with a p-value threshold of 0.0001 and a deviation parameter of 2.5. These conservative parameters were chosen to minimize the potential influence of false-positive reassortment events on the analysis that may appear due to sequencing errors or inconsistencies in substitution rates of genes.

Similarly, we performed the reassortment analysis on the H5Nx clade 2.3.4.4b strains. We determined the clade membership using the Nextclade H5Nx classifier [2] and split out a major monophyletic clade of H5N1 strains that appeared in 2020. That way, we analyzed the background H5Nx 2.3.4.4b reassortment rates and patterns separately from the 2020-2023 H5N1 2.3.4.4b clade.

## Acknowledgments

We gratefully acknowledge pork producers, swine veterinarians, and laboratories for participating in the USDA Influenza A Virus in Swine Surveillance System and publicly sharing sequences. We also gratefully acknowledge all data contributors, i.e., the Authors and their Originating laboratories responsible for obtaining the specimens, and their Submitting laboratories for generating the genetic sequence and metadata and sharing via the GISAID Initiative, on which components of this research is based. This work was supported in part by the USDA-ARS (ARS project number 5030-32000-231-000D); USDA-APHIS (ARS project number 5030-32000-231-080-I); the National Institute of Allergy and Infectious Diseases, National Institutes of Health, Department of Health and Human Services (Contract No. 75N93021C00015); the Centers for Disease Control and Prevention (contract numbers 21FED2100395IPD, 24FED2400250IPC); and the SCINet project of the USDA-ARS (ARS project number 0500-00093-001-00-D). The funders had no role in study design, data collection and interpretation, or the decision to submit the work for publication. Mention of trade names or commercial products in this article is solely for the purpose of providing specific information and does not imply recommendation or endorsement by the USDA or CDC. USDA is an equal opportunity provider and employer.

## Supplemental Materials

**Figure S1:**
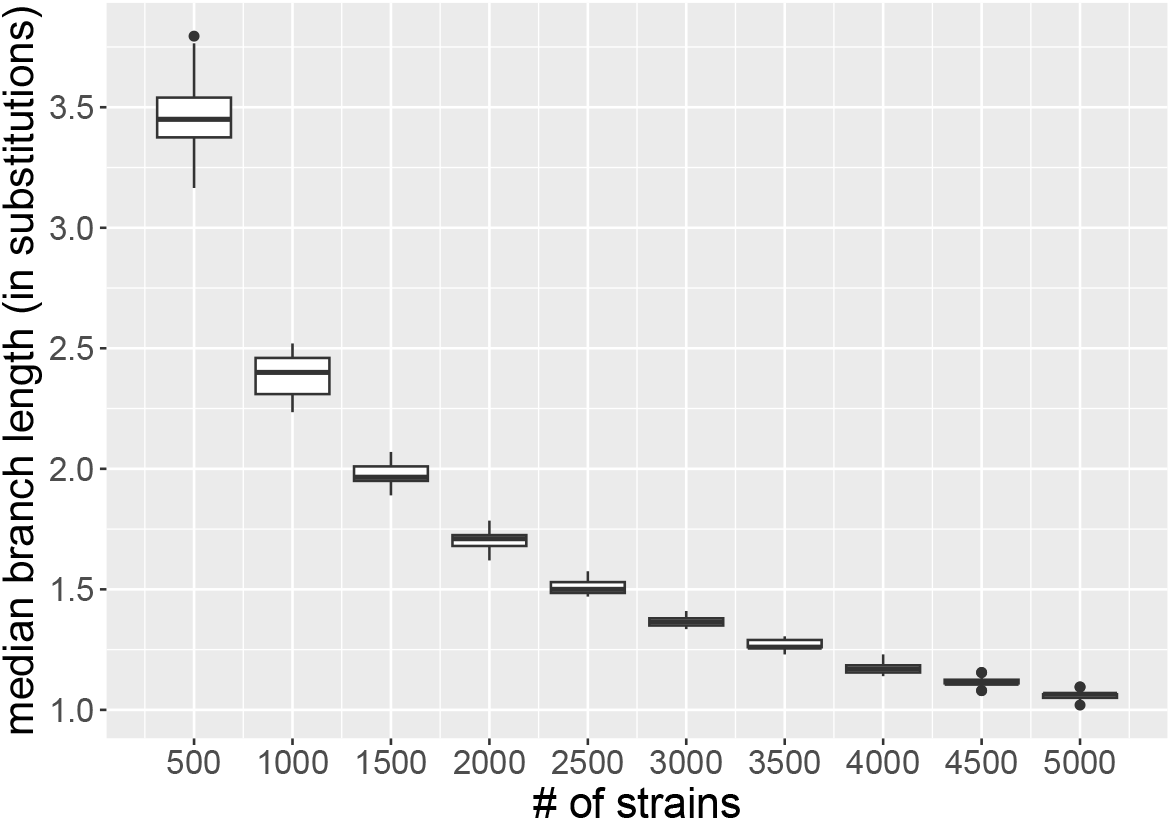
The sampling density (median branch length) of the datasets in the simulation study. The median branch length, measured in the number of substitutions per branch, provides a standardized metric comparable across different datasets, and it decreases as the sampling density increases. In simulations, we show that TreeSort is very accurate on datasets with the median branch length below 2.

**Figure S2:**
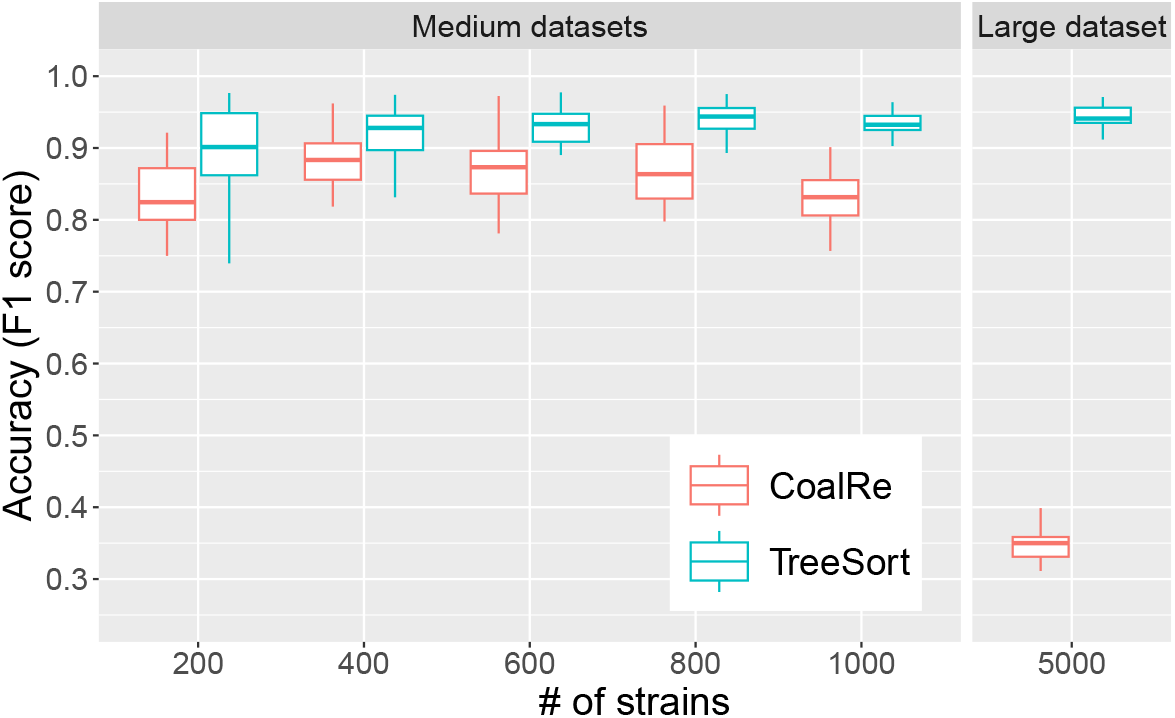
The accuracy comparison between TreeSort and CoalRe by Muller et al. on simulated data. The accuracy is measured in terms of the F1 score that combines the precision and recall metrics [38].

**Figure S3:**
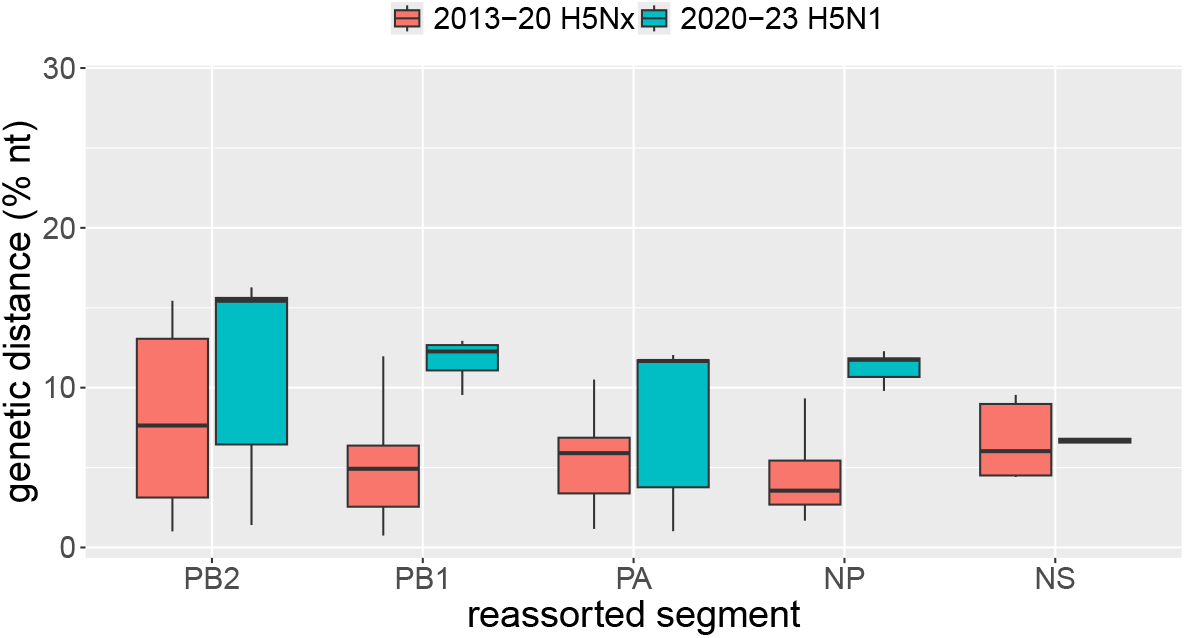
The distance between the genes swapped by reassortment events in two different periods of H5Nx 2.3.4.4b evolution. For example, a genetic distance of 10% for a PB2 gene implies that a single reassortment event occurred where the novel PB2 gene was 10% different from the original PB2. Overall, the H5N1 2.3.4.4b virus in the 2020-2023 period underwent more significant reassortment events than H5Nx 2.3.4.4b in the 2010-2020 period. We omitted NA and MP segments from the comparison due to the low number of reassortment events with these segments in the 2020-2023 period.

**Table S1:** Substitution rates per site per year for H5Nx clade 2.3.4.4b viruses in two different periods (2010-2020 and 2020-2023) estimated by TreeTime v.0.11.3.

| Segment | 2010-2020 subst. rate | 2020-2023 subst. rate |
| --- | --- | --- |
| PB2 | 0.0020 | 0.0035 |
| PB1 | 0.0024 | 0.0039 |
| PA | 0.0023 | 0.0055 |
| HA | 0.0045 | 0.0055 |
| NP | 0.0018 | 0.0040 |
| NA | 0.0027 | 0.0063 |
| MP | 0.0027 | 0.0061 |
| NS | 0.0050 | 0.0067 |

**Figure S4:**
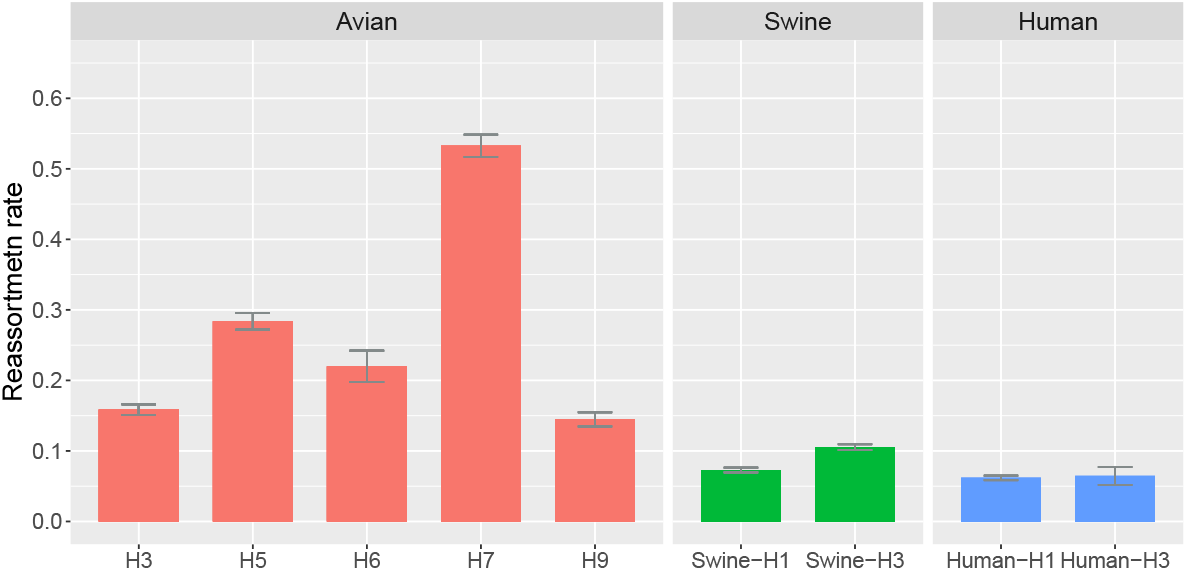
Reassortment rate estimates via the 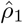 estimator for major avian, swine, and human subtypes. The reassortment patterns are similar to those shown in Fig. 2A with the 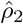 estimator.

## Footnotes

∗ Here and throughout the Results, we refer to the 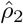 estimate of the reassortment rate (see Methods).

